# Ultra-fast genome-wide inference of pairwise coalescence times

**DOI:** 10.1101/2023.01.06.522935

**Authors:** Regev Schweiger, Richard Durbin

## Abstract

The pairwise sequentially Markovian coalescent (PSMC) algorithm and its extensions infer the coalescence time of two homologous chromosomes at each genomic position. This inference is utilized in reconstructing demographic histories, detecting selection signatures, genome-wide association studies, constructing ancestral recombination graphs and more. Inference of coalescence times between each pair of haplotypes in a large dataset is of great interest, as they may provide rich information about the population structure and history of the sample.

We introduce a new method, *Gamma-SMC*, which is *>*14 times faster than current methods. To obtain this speed up, we represent the posterior coalescence time distributions succinctly as a Gamma distribution with just two parameters; while in PSMC and its extensions, these are held as a vector over discrete intervals of time. Thus, Gamma-SMC has constant time complexity per site, without dependence on a number of discrete time states. Additionally, due to this continuous representation, our method is able to infer times spanning many orders of magnitude, and as such is robust to parameter misspecification. We describe how this approach works, illustrate its performance on simulated and real data, and use it to study recent positive selection in the 1000 Genomes Project dataset.

## 1 Introduction

Each of an individual’s two alleles, at each genomic position, is inherited from a chain of ancestors going back in time. These two chains must eventually coalesce at an identical common ancestor. The length of these chains, measured by the (scaled) number of generations until coalescence, is called the *coalescence time*, or the *time to the most recent common ancestor* (TMRCA). Coalescence times reflect the genealogical and genetic history of a sample, and therefore have been used in many population and statistical genetic analyses, such as reconstructing demographic histories [15, 25, 27, 28], detecting selection signatures [11, 21, 22], genome-wide association studies [20, 29], genotype imputation [29] and more.

As genomic datasets grow larger, inference of coalescence times between each pair of haplotypes in a large dataset is of great interest, as they may provide rich information about the population structure and history of the sample in resolution impossible to attain otherwise. Moreover, they may further be used to construct an ancestral recombination graph (ARG) [8] - a sequence of local genealogies that documents the genealogical history of each locus across the sample.

Coalescence times along two homologous chromosomes can be well approximated with a Markov chain using the sequentially Markovian coalescent (SMC) framework [16, 19]. This framework underlies the PSMC [15] method, which estimates a posterior distribution of the TMRCAs, as well as infers a history of change in the effective population size over time. While PSMC was designed to work on a single pair of sequences, follow-up methods have been developed to scale to increasingly larger sample sizes (for a review, see [17]). The Multiple Sequentially Markovian Coalescent (MSMC) [25] analyses the genomes of multiple individuals, focusing on the first coalescence between any two individuals; but is computationally prohibitive, and can only be applied to up to 8 samples. MSMC2 [28] employs a composite likelihood approach instead, but is still limited in sample size. The diCal method [10] allows decoding coalescent times in linear time with respect to the state space of an HMM, instead of the quadratic complexity in PSMC, MSMC and MSMC2. The SMC++ method [27] allows incorporating allele frequencies from hundreds of unphased genomes to decode coalescence times more accurately for a focal pair of haplotypes. Finally, the recent ASMC method [22] incorporates computational and algorithmic improvements that reduce running times by 2-4 orders of magnitude, and extends the approach to work on genotyping arrays. Despite these advances, further scaling TMRCA inference is still desirable.

To facilitate such large-scale analyses, we introduce a new method, Gamma-SMC, which is 14-20 times faster than current methods. To obtain this speed up, we represent the posterior coalescence time distributions succinctly as a gamma distribution with just two parameters, rather than as a vector over discrete intervals of time as in PSMC and its extensions. Thus, Gamma-SMC has constant time complexity per site, without dependence on a number of discrete time states. Additionally, due to this continuous representation, our method is also able to infer times spanning several orders of magnitude, and as such is robust to parameter misspecification. We describe how this approach works and illustrate its performance on simulated and real data. Finally, we use the estimated coalescence times to scan for sites under recent positive selection in the human genome.

## 2 Results

### 2.1 Overview of the Gamma-SMC method

Like its predecessors [15, 22, 26, 27], Gamma-SMC is based on an HMM, whose transition probabilities are derived from the SMC framework. In SMC-based methods, the hidden states along the genome correspond to pairwise TM-RCAs, and the emissions are the observed diploid sequence. However, standard HMMs require a discrete hidden state space, while coalescence time is naturally continuous. Therefore, current SMC-based methods approximate the full model by discretizing time into a number of intervals (e.g., *T* = 20, 32 or 64) and so have at least *O*(*T*) computational complexity per step.

#### Avoiding time discretization with a Gamma approximation

We now describe how Gamma-SMC avoids time discretization. We focus here on the forward algorithm of the HMM, but similar arguments hold for the backward algorithm, and for combining both forward and backward passes (Methods). The forward algorithm iteratively updates the posterior distribution of the hidden state, conditional on the observations seen so far [6, 23]. That is, it measures and updates our uncertainty of the coalescence time at a locus, given the sequence up until that locus. Instead of holding this distribution as a vector as in current methods, we observed, as others have done [4], that the SMC posteriors are very well approximated by a gamma distribution. If we constrain the posterior to be a member of the family *I’* (*α, β*), then the hidden state is described by only two parameters (*α* and *β*) per step. Analogues of all standard HMM algorithms have been derived for CS-HMMs [3] and in principle this leads to *O*(1) time-complexity per step, compared to *O*(*T*) or even *O*(*T* ^2^) by current methods.

The motivation for using the gamma distribution is threefold. First, the gamma distribution is a rich family, well suited to model positive, unimodal and continuous distributions, with only two parameters. Second, the gamma distribution is conjugate to the Poisson distribution, which underlies the emission probability in the HMM. This means that, without recombination, and with the natural exponential prior on coalescence time, the forward densities should be exactly gamma. It is therefore reasonable that with a small recombination probability per step, the Gamma shape may still approximate the true density well. Finally, an empirical study of forward densities of PSMC shows a Gamma distribution approximation often well approximates the true density [4].

To support HMM operations, we need to describe how to update the forward density from one step to the next. As mentioned above, following recombination the forward density in the next step need not be gamma, even if it was gamma at the previous step. Fortunately, for small recombination rates, consecutive forward densities will be very similar, so we can likely approximate the next step with gamma as well. The forward algorithm alternates between two steps: Multiplying the current forward density by the transition density, to account for additional uncertainty arising from a possible recombination event; and updating the new density using the emission probability, to account for the additional information from the new observation. We explain each in turn.

For the emission step, we can exploit the conjugacy between Poisson and gamma to derive a simple update rule for the *α, β* parameters. Denote the *i* + 1-th observation by *y ∈* {0, 1}, indicating if this is a heterozygous (het) site, where the sequences dier, or a homozygous (hom) site, identical in both sequences. If the posterior density of the TMRCA at time *i* + 1, conditional on the first *i* observations, is *Γ* (*α,β*), then the forward density in step *i*+1 is simply *Γ* (*α* + *y, β* + 2*θ*), where is the scaled mutation rate.

For the transition step, we derived a closed-form expression of the coalescence time at step *i*+1, given that the forward density at step *i* is gamma (Supplement). This expression is not in the form of a gamma distribution, but for a small recombination rate, is expected to be close. Therefore, for each possible *Γ* (*α,β*) distribution, we find a new approximate subsequent *Γ* (*α*′, *β*′) distribution. The mapping ℱ: (*α,β*) → (*α*′, *β*′) may be viewed as a flow field, or a two-dimensional vector field, describing a dynamical system that develops along the genome. We evaluated this flow field over a log-spaced grid (Methods), and use bilinear interpolation for off-grid values. Importantly, while calculating this flow field is computationally expensive, it needs only be calculated once.

#### Locus skipping

As in other SMC methods, we preprocess the genome into segments. Each segment ends with either a het site, a site defined by the user as a target for inference, or when a maximal length (e.g. 10kbp) is reached. Within each segment are only hom sites (for discussion of missing sites, see Methods). We calculate in advance the result of repeatedly applying transition and emission steps with hom sites, up until maximal segment length, and cache these results. Then the forward pass can effectively skip stretches of hom sites, and be evaluated only at segment ends. This allows for additional substantial speed up.

#### Coding

We implemented our method in C++, using vectorized SIMD operations. A preliminary version used to generate the results in this paper is available from https://www.github.com/regevs/gamma_smc.

### 2.2. Benchmarks

#### Accuracy

We conducted a simulation study to evaluate the accuracy of Gamma-SMC, compared to current SMC-based methods. We simulated a whole genome using realistic population genetic parameters (Methods), and used ASMC-seq [22], MSMC2 [28] and Gamma-SMC to infer the posterior mean of the coalescence times. When comparing the true TMRCA and its inferred posterior mean (Figure 1), we conclude that all methods have similar inference accuracy, with ASMC-seq more accurate and Gamma-SMC less accurate. This is further confirmed by measuring Pearson’s correlation *r*^2^ (ASMC-seq = 0.85, MSMC2 = 0.82, Gamma-SMC = 0.80) and the mean absolute error (in generations, ASMC-seq = 8,687.5, MSMC2 = 10,349.4, Gamma-SMC = 10,547.7) between the true and inferred TMRCA. Although Gamma-SMC performance here is lower than that of the discretised HMM algorithms, as we will see below, it is good enough for further inference.

**Fig. 1:**
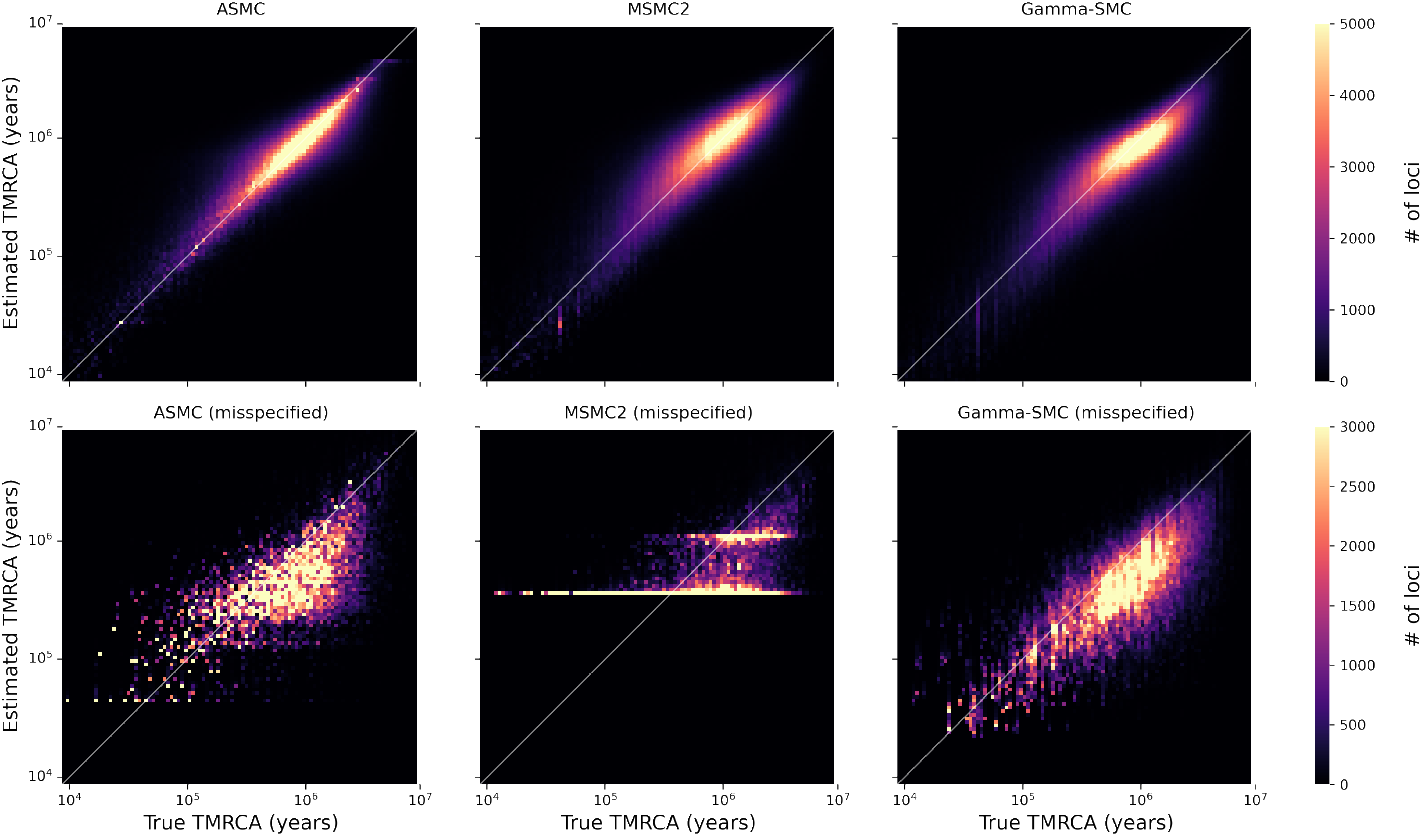
Inference accuracy of Gamma-SMC vs. MSMC2 and ASMC-seq. A comparison of the true TMRCA vs. the estimated TMRCA (posterior mean) across a simulated genome. Shown for ASMC-seq (top left), MSMC2 (top middle) vs. Gamma-SMC (top right) when the correct population genetic parameters are given; and when the TMRCAs are × 100 smaller than expected (MSMC2, bottom left; Gamma-SMC, bottom right).

We next analysed the distribution of the posterior mean TMRCA estimator across ASMC-seq, MSMC2 and Gamma-SMC (Figure 2a). We note that, while all methods tend to regress to the mean, overestimating low TMRCA values and underestimating high TMRCA values, mean Gamma-SMC estimates are lower than mean ASMC-seq and MSMC2 estimates in each bin of true TMRCA, suggesting there might be a slight consistent bias which could be corrected.

**Fig. 2:**
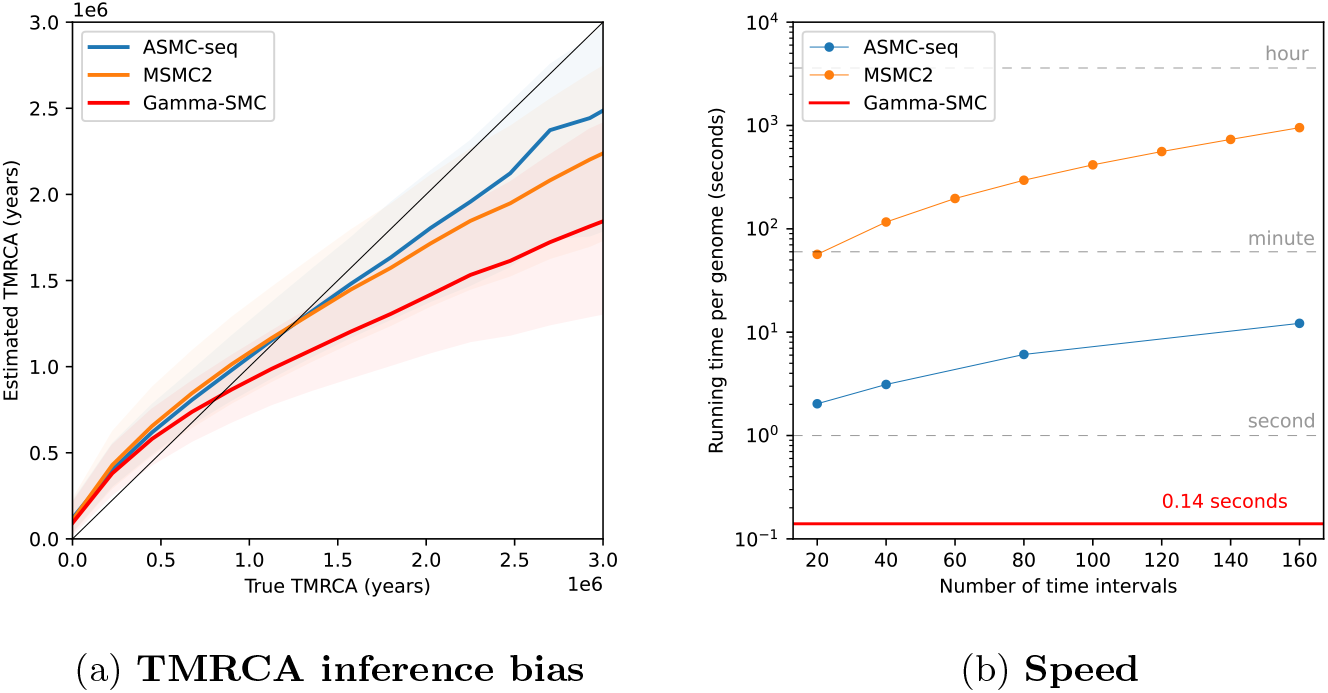
(a) A study of TMRCA inference bias in Gamma-SMC vs. MSMC2 and ASMC-seq. The mean inferred TMRCA is shown as a function of the true TM-RCA (confidence band, 25%-75% quantiles). (b) Running time of Gamma-SMC vs. MSMC2 and ASMC-seq, for a whole genome (3.2Gb).

#### Robustness to scale

Another drawback of the time discretisation employed by current methods is that inference resolution is determined by the discretisation scheme chosen, so that it is complex if not impossible to reliably distinguish between TMRCAs in the same time interval. In particular, inference will be poor if the true TMRCA is very small - this may happen, for example, in a long run of homozygosity, or with parameter misspecification.

To this end, we simulated a diploid sequence and used ASMC-seq, MSMC2 and Gamma-SMC to estimate TMRCAs, but used in simulation an effective population size of 150, effectively decreasing by a factor of 100 the true underlying TMRCAs relative to the tuning of the algorithms. When such small TMRCAs are estimated by MSMC2, the interval boundaries make inference coarse and imprecise, while with Gamma-SMC inference is much less affected (Figure 1). ASMC-seq inference is better than MSMC2, but worse than Gamma-SMC (*r*^2^ (ASMC-seq = 0.55, MSMC2 = 0.41, Gamma-SMC = 0.67) We note that the effective range in Gamma-SMC is determined only by the grid values the flow field was pre-calculated on, such that arbitrarily small or large TMRCAs can be inferred if needed.

#### Demographic model

Gamma-SMC assumes an unrealistic constant-size population demographic model its flow field construction. To test the effect of this assumption, we simulated a human genome using the out-of-africa model of [9], as implemented by *stdpopsim* [2], and inferred TMRCAs using Gamma-SMC (see Supplement). We observed that: (i) Gamma-SMC displays a consistent upward bias throughout the entire range; (ii) *r*^2^ for the out-of-africa simulation is slightly better than for the constant-size simulation (*r*^2^ = 0.85 vs 0.8); (iii) similarly, mean absolute error is slightly lower for out-of-africa (8885.6 generations) than for the constant model (10547.7 generations). We conclude that Gamma-SMC is still helpful even in face of some demographic model mis-specification, although care must be taken when using absolute quantities, rather than relative estimates.

#### Speed

We next evaluated the running time of Gamma-SMC, and compared it to the state-of-the-art method, ASMC-seq, as well as to MSMC2. The running time of Gamma-SMC in practice is affected by several factors, such as the number of segregating sites, patterns of missing data, and more. For a fair comparison, we used the example provided in the ASMC git repository - 5,086 segregating sites within 30Mbp, across 150 diploids (300 haplotypes, 44,850 haplotype pairs), with no missing sites.

We observed that the running time of Gamma-SMC is at least ×14 faster than ASMC-seq (Figure 2b), depending on the number of discrete TMRCA intervals used in ASMC-seq. For example, analysis of all pairs using *T* = 69 discrete time intervals, as is the smallest pre-computed default provided by ASMC-seq, required 29 minutes for a single processor for ASMC-seq, compared to ∼1 minute for Gamma-SMC. Extrapolated to a genome size of 3.2Gbp, Gamma-SMC can process a whole genome in ∼0.14 seconds.

### 2.3 Application to 1000 Genomes Project data

We applied Gamma-SMC to 503 individuals of 5 populations of European ancestry from the 1000 Genomes Project dataset [1] (Phase 3). We inferred the posterior distribution of the TMRCAs, for each pair of the 1,006 haplotypes, at evenly-spaced intervals of 1,000 basepairs across all chromosomes. Excluding data loading and writing results, this took a total of 120.5 CPU hours for 505,515 haplotype pairs, computed in parallel across 7,150 jobs. (At 0.85 sec per pair, this is slower than the value given above, essentially because of masking and pre- and post-processing.)

We illustrate the results by showing an example of inference at a single position (Figure 3). We show a heatmap of the posterior mean TMRCA between each pair of haplotypes. The order of haplotypes is given by UPGMA hierarchical clustering. We note the clear hierarchical structure, corresponding to the coalescent tree at this position.

**Fig. 3:**
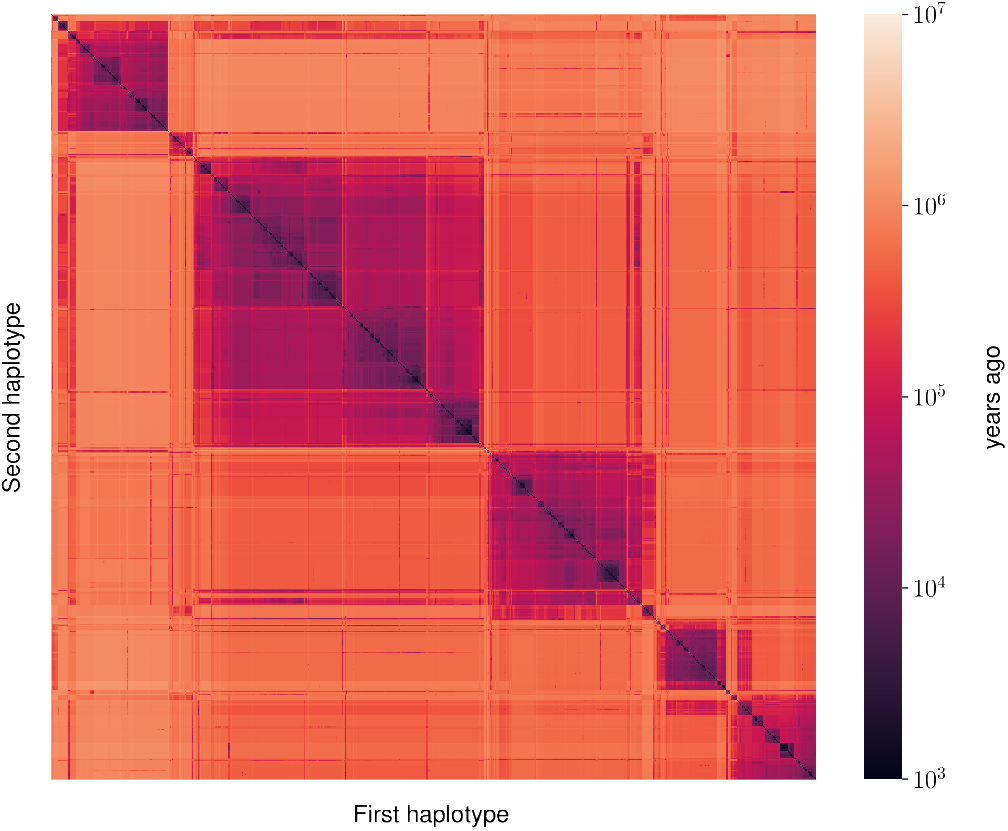
Coalescence times in 1000 Genomes Project data. Pairwise co-alescence time for all haplotype pairs of European individuals from the 1000 Genomes Project data, at chr2:500000. Coalescence times are estimated by taking posterior means, measured in years ago, assuming generation time of 30 years and effective population size of 15,000.

To further demonstrate the utility of Gamma-SMC, we used it to detect loci under recent positive selection. Positive selection at a locus causes a rapid rise in the frequency of the beneficial allele. As such, haplotypes with the beneficial allele will coalesce much faster than for neutral sites. When selection is recent, this causes an enrichment for recent coalescent times in the distribution of pairwise TMRCAs at positively selected loci. This skew can be used as a basis for a test for selection [11, 22].

We performed a selection scan by looking for genomic regions enriched for recent coalescent times (Figure 4a); specifically, regions of size 100kbps where a large percentage of pairwise coalescences are estimated to have occurred in the last 4,500 years. We recover approximately 20 candidate loci, including for example LCT [18], the gene encoding the lactase enzyme, and the HLA region, both known to have been under recent positive selection in Europeans (Figure 4b). The peak at chromosome 12 falls inside HECTD4, which has recently been found to be under selection in Eurasians [13], though we note that the credible interval contains other genes.

**Fig. 4:**
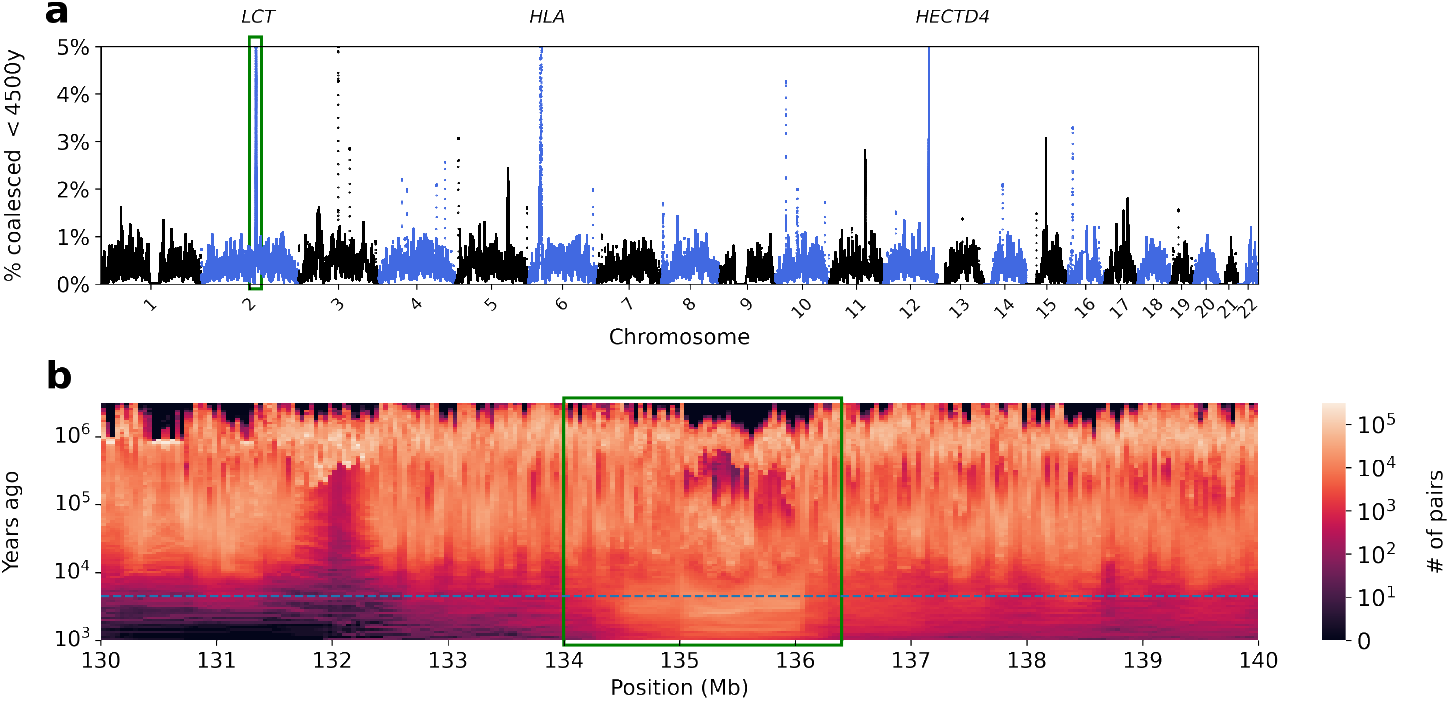
Scanning for recent positive selection. **(a)** We searched for 100kbp regions enriched for pairwise coalescence times more recent than 4,500 years ago. **(b)** A focus on the locus of the LCT gene (green rectangle), showing an enrichment of coalescence time in recent years (dashed line, 4,500 years ago, assuming generation time of 30 years and effective population size of 15,000).

## 3 Discussion

We present Gamma-SMC, a method for efficiently inferring coalescence times from sequencing data. We showed that Gamma-SMC is at least an order of magnitude faster than current methods, and that it can infer very small or very large TMRCAs accurately, due to its continuous state space. Finally, we applied Gamma-SMC to 505,515 pairs of haploid genomes from the 1000 Genomes Project data, and detected evidence for recent positive selection at multiple sites across the genome.

We note several limitations of Gamma-SMC. As with other SMC-based methods, good quality phasing is required to apply Gamma-SMC across individuals. While possible for large datasets, it may be harder to obtain for smaller ones. In addition, while the accuracy of Gamma-SMC is comparable to current methods, it could be further improved. Directions include incorporating genetic maps for inference; using more accurate demographic models when building the flow field; incorporating a post-hoc TMRCA normalization step [29]; and using the conditioned sample frequency spectrum [22, 27], which utilizes allele frequencies in inference. In terms of speed, GPUs may offer additional acceleration. Finally, Gamma-SMC currently works only on sequencing data. Extending Gamma-SMC to use the conditional Simonsen-Churchill model [12] would enable it to operate on SNP genotyping array data, and is a promising direction for future research.

One exciting potential application of Gamma-SMC is in constructing an ARG. Indeed, the clear hierarchical structure evidenced in the pairwise posterior TMRCAs (Figure 3) reflects the genealogical tree structure at a site which, when inferred along the genome, gives rise to the tree sequence representation of an ARG [29]. Gamma-SMC may improve branch placement and timing in the weaving step of ARG building methods such as ARGweaver [24] and ARG-Needle [29], as the continuous nature of the posterior distributions of coalescence times in Gamma-SMC may help resolve ambiguities present in discrete time approximations.

A disadvantage of current scalable ARG building methods is that they only output a single ARG. Further focusing on a particular locus, there is interest in sampling from the posterior distribution of trees at a single position, as this allows quantifying the uncertainty in both dating of coalescence events and the topology of the tree. Gamma-SMC gives a pairwise coalescence posterior distribution, which supports sampling, and also is fast enough to allow iterated application to inferred ancestral sequences arising from initial, accepted coalescences.

Previous methods also infer the history of population size of the sample, but inference is limited in recent times due to sparsity of recent coalescence events. However, the number of pairs in a population sample grows quadratically with its size, so that for a large sample, many more recent coalescence events are expected. Utilizing this for demography inference is another promising avenue of research.

## 4 Acknowledgments

We thank Trevor Cousins and Aylwyn Scally for helpful discussions, and Fredrik Johansson for support with the Arb library. R.S. is supported by an EMBO Postdoctoral Fellowship (ALTF 783-2020). This research was funded in part by the Wellcome Trust grant WT207492. For the purpose of open access, the author has applied a CC BY public copyright licence to any Author Accepted Manuscript version arising from this submission. R.S. is a paid consultant to MyHeritage Ltd. and Eleven Therapeutics.

## 5 Methods

### 5.1 Gamma-SMC

We outline the structure of the full Gamma-SMC algorithm, while deferring most mathematical derivations to the Supplementary Methods.

#### Flow field

Define ℱ to be the flow field mapping from (*α,β*) to (*α*′, *β*′), where *Γ* (*α*′, *β*′) is a gamma approximation of the density of the compound distribution given by marginalizing over the density of *Γ* (*α,β*) according to the SMC’ transition probabilities. We describe how to evaluate the compound distribution analytically in the case of constant population size, and hence how we fit ℱ (*α, β*), in A.4.

Prior to any analysis, and independently of any parameters, we evaluate over a 2-dimensional grid of values. In practice, instead of *α, β*, we evaluate over a grid of *l*_*μ*_ := log_10_(*μ*) and *l*_*C*_ := log_10_(*CV*), where *μ* = *α/β* and 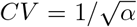 are the mean and coefficient of variation of *Γ* (*α,β*), respectively. The grid is defined by 51 *l*_*μ*_ values equally spaced between −5 and 2 (that is, log-spaced between *μ* = 10^−5^ and 10^2^), and 50 *l*_*C*_ values equally spaced between −2 and 0 (CV between 10^−2^ and 1). Units here are in coalescent time. We exclude CV*>*1 as this results in an infinite gamma density at *x* = 0. We make extensive use of the Arb library [14].

#### Segmentation and caching

Segregating sites are sparse, between long stretches of homozygosity. This can be used for “locus skipping” - performing a single step of the forward algorithm that accounts for a stretch of identical observation, based on a cached calculation. To this end, we calculate, for each element of the flow field grid, and for each number of steps from 1 to a maximal cache size, the result of repeatedly applying the flow field, and incorporating a hom observation. We calculate a similar set of values for stretches of missing values. The results (effectively two new flow fields, per number of steps) are cached and used in the forward pass.

#### Input and output

Gamma-SMC works on a VCF file. In addition, it optionally accepts a list of BED files, one per sample in the VCF; each BED file describes a mask of genomic intervals outside which alleles are considered missing. The user also describes at which sites to infer the TMRCA posterior - either at heterozygous sites, across a grid of evenly-spaced positions, or both. The input is then segmented as described in the main text.

Within each segment, we need to calculate how many basepairs are homozygous, and how many are missing. For each pair of samples and for each segment, we calculate the total length of the intersection of corresponding two masks and the segment. Then, we treat the segment as if it is made of a stretch of missing observations followed by a stretch of homozygous observations. The user must also provide the scaled mutation rate *θ* and scaled recombination rate *r*.

We output the final posterior gamma approximation, for each pair of haplotypes, at each output position.

#### Forward pass

For each pair of haplotypes, we perform the forward pass: For each segment, we begin from the (*l*_*μ*_, *l*_*C*_) parameters describing the gamma approximation to the forward density at the end of the previous segment (or (0, 0) for the first segment). We look up in the cache the result of applying of the flow field on a stretch of missing values of the length we observe, and get new a (*l*_*μ*_, *l*_*C*_) pair; we then look up in the cache the result of applying the flow field on a stretch of hom value of the length we see; finally we incorporate the observation at the end of the segment, as explained in the Supplement.

When looking up a pair of values in a flow field, if one of the (*l*_*μ*_, *l*_*C*_) values lies beyond the grid range, it is clipped back to the grid boundaries. Otherwise, they are interpolated (bilinear interpolation) using the 4 nearest neighbours.

We record the result of the forward pass at the output positions - that is, for each site required by the user for TMRCA inference, we record the (*l*_*μ*_, *l*_*C*_) we obtained from the forward pass, describing the gamma approximation to the forward density at that site.

#### Backward pass

Instead of running the CS-HMM equivalent of the backward algorithm, instead we re-formulate the backward pass in terms of applying the forward pass on the reverse sequence, with an additional transition through the flow field (Supplement). We again record the gamma approximation of the backward pass at the output position. Finally, we combine the forward and backward densities - if *Γ* (*α, β*) approximates the forward density at a position, and *Γ* (*α*′, *β*′) the backward, then the combined posterior density will be *Γ* (*α* + *α*′ *-* 1, *β* + *β*′ *-* 1) (Supplement).

### 5.2 Simulation study

We simulated 22 human chromosomes, using msprime v1.0.2 [5]. We used an effective population size of *N*_*e*_ = 15, 000 (30,000 haploids), mutation rate *μ* = 1.25 · 10^−8^, and a human recombination map for each chromosome. We used ASMC v1.2 in sequencing mode.

