## Supplementary Information for "Ultra-fast genome-wide inference of pairwise coalescence times"

### A Supplementary Information

#### A.1 Out-of-africa model

Figures A.1 and A.2 show an analysis analogous to Figures 1 and 2, on the out-of-africa model of [9].

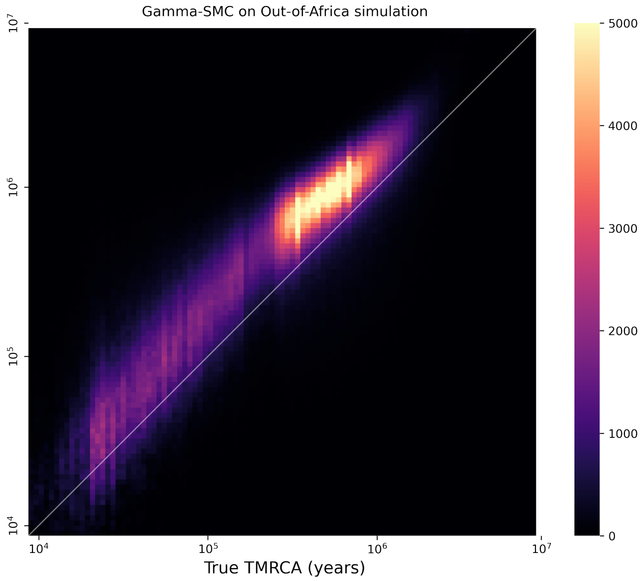


Fig. A.1: **Inference accuracy of Gamma-SMC for out-of-africa model** A comparison of the true TMRCA vs. the estimated TMRCA (posterior mean) across a genome according to an out-of-africa model.

#### A.2 The Sequentially Markovian Coalescent (SMC) model

We begin by describing the SMC. For a detailed overview, see e.g, the Supplement of [28].

*Hidden states and observations.* An observation $Y_i$ ($i = 1, \dots, N$, with $N$ the sequence length) at location $i$ in the genome for a pair of haplotypes, can take a value from $Y_i \in \{-1, 0, 1\}$, where -1 denotes missing data in either of the two haplotypes (not called, or masked out; see below), 0 denotes a site where both haplotypes have the same allele (i.e. a homozygous genotype in case of a single diploid genome), 1 denotes a mismatch between the alleles of the two haplotypes (i.e. a heterozygote genotype in case of a single diploid genome). The hidden states $\{X_i\}_{i=1}^N$ are the TMRCAs between the two alleles at position $i$.

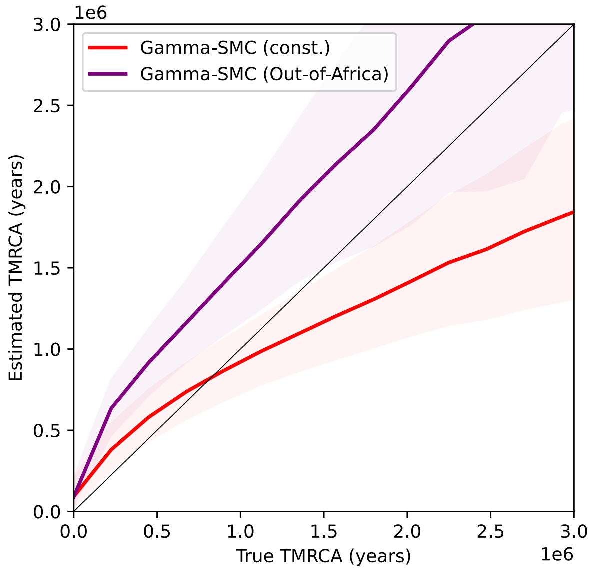


Fig. A.2: A study of TMRCA inference bias in Gamma-SMC, for genomes simulated with either a constant population size or an out-of-africa model. The mean inferred TMRCA is shown as a function of the true TMRCA (confidence band, 25%-75% quantiles).

*Emission probability function.* The emission probabilities for a given coalescence time $s$ follow a Poisson distribution, with rate $\lambda = 2\mu s$. We assume $\mu \ll 1$, so that the probability of recurrent mutations is negligible:

$$\begin{aligned}\Pr(Y_i = -1 \mid X_i = s) &= 1 \\ \Pr(Y_i = 0 \mid X_i = s) &= e^{-2\mu s} \\ \Pr(Y_i = 1 \mid X_i = s) &= 1 - e^{-2\mu s}\end{aligned}$$

*Transition density function.* The transition probability is derived from the SMC model by McVean and Cardin [19], as amended in the SMC' model by Marjoram and Wall [16]. A recombination event, which takes place at position $i$, detaches the chain of ancestors, which then coalesces back, either onto itself or the other branch, at time $t$. Formally, define:

$$q(t|s) := \Pr(X_{i+1} = t | X_i = s, R_i)$$

where $R_i$ denotes the event of having a recombination event between position $i$ and $i + 1$. We assume a constant population size throughout. In this case, this is given by [7], Eq (6):

$$q(t|s) = \begin{cases} \frac{2t+e^{-2t}-1}{4t} \cdot \delta(t-s) & t = s, \\ \frac{1-e^{-2t}}{2s} & t < s, \\ \frac{e^{-(t-s)}-e^{-(t+s)}}{2s} & t > s. \end{cases}$$

Also conditioning on the event of recombination, we get

$$\begin{aligned} p(t|s) :=& \Pr(X_{i+1} = t|X_i = s) \\ =& \Pr(X_{i+1} = t|X_i = s, R_i) \cdot P(R_i|X_i = s) + \delta(t-s) \cdot P(\bar{R}_i|X_i = s) \end{aligned}$$

Where, again we assume $\rho \ll 1$, so we neglect the event of more than one recombination.

##### A.3 Posterior density in continuous-state HMMs (CS-HMMs)

*The forward algorithm.* The forward algorithm in the context of a CS-HMMs [3] is an iterative procedure which tracks the probability density (*forward density*) of a hidden state at a position $i$, given the observations until that position. We use the scaled variant [6]. The forward densities are defined by:

$$\hat{\alpha}(x_i) = \Pr(X_i = x_i|\mathbf{Y}_{1:i} = \mathbf{y}_{1:i})$$

with the recursion

$$\hat{\alpha}(x_i) = \frac{1}{c_i} \cdot \Pr(y_i|x_i) \int_0^\infty \Pr(x_i|x_{i-1})\hat{\alpha}(x_{i-1})dx_{i-1}.$$

with $c_i$ a scaling factor that assures we get a normalized density function. We begin with an exponential prior $\hat{\alpha}(x_0) \sim \Gamma(1,1)$, as this is the stationary distribution of the SMC'.

*Posterior state density.* We use a slightly different alternative to the backward algorithm, which is easier to work with. Denote by $\overleftarrow{\alpha}(x_i)$ the result of running the scaled forward algorithm on the reversed sequence; that is,

$$\overleftarrow{\alpha}(x_{i+1}) = \Pr(X_{i+1} = x_{i+1}|\mathbf{Y}_{i+1:N} = \mathbf{y}_{i+1:N})$$

Taking this one step back, we get:

$$\Pr(X_i = x_i|\mathbf{Y}_{i+1:N} = \mathbf{y}_{i+1:N}) = \int_0^\infty \Pr(x_i|x_{i+1})\overleftarrow{\alpha}(x_{i+1})dx_{i+1}$$

Then, we have:

$$\begin{aligned}
\Pr(x_i|\mathbf{y}_{1:N}) &= \frac{\Pr(\mathbf{y}_{1:N}|x_i)\Pr(x_i)}{\Pr(\mathbf{y}_{1:N})} \\
&= \frac{\Pr(\mathbf{y}_{i+1:N}|x_i)\Pr(\mathbf{y}_{1:i}|x_i)\Pr(x_i)}{\Pr(\mathbf{y}_{1:N})} \\
&= \frac{\Pr(\mathbf{y}_{i+1:N}|x_i)\Pr(x_i|\mathbf{y}_{1:i})\Pr(\mathbf{y}_{1:i})}{\Pr(\mathbf{y}_{1:N})} \\
&= \hat{\alpha}(x_i)\cdot\frac{\Pr(\mathbf{y}_{i+1:N}|x_i)\Pr(\mathbf{y}_{1:i})}{\Pr(\mathbf{y}_{1:N})} \\
&= \hat{\alpha}(x_i)\cdot\frac{\Pr(x_i|\mathbf{y}_{i+1:N})\Pr(\mathbf{y}_{i+1:N})\Pr(\mathbf{y}_{1:i})}{\Pr(x_i)\Pr(\mathbf{y}_{1:N})} \\
&= \hat{\alpha}(x_i)\cdot\int_0^\infty \Pr(x_i|x_{i+1})\overleftarrow{\hat{\alpha}}(x_{i+1})dx_{i+1}\cdot\frac{1}{\Pr(x_i)}\cdot\underbrace{\frac{\Pr(\mathbf{y}_{i+1:N})\Pr(\mathbf{y}_{1:i})}{\Pr(\mathbf{y}_{1:N})}}_{Constant}
\end{aligned}$$

Namely, the posterior density of the TMRCA at a position $i$ can be obtained by: (i) Running the forward algorithm until step $i$; (ii) Running the forward algorithm on the reversed sequence until step $i+1$; (iii) evaluating the integral one step back, to step $i$; (iv) divide by the prior density of $x_i$; (v) scale to obtain a legal density function.

##### A.4 Gamma approximation

**Forward algorithm.** As mentioned above, assuming a standard coalescent, with constant population size, the coalescence time has an exponential prior; that is, $\Pr(x_i) \sim Exp(1) = \Gamma(1,1)$. Now, assume $\hat{\alpha}(x_i) \sim \Gamma(\alpha,\beta)$. We describe how to approximate $\hat{\alpha}(x_{i+1})$.

*Transition step.* We give a closed-form expression for the coalescence time at the next step. We first take a linear approximation, assuming the recombination rate is small:

$$\Pr(R_i|X_i = s) \approx 2\rho s$$

Denote by $f_{\alpha,\beta}(x)$ the pdf of the gamma function $\Gamma(\alpha,\beta)$. Define

$$p_{\alpha,\beta}(t) := \Pr(X_{i+1} = t|X_i) \quad \text{s.t.} \quad X_i \sim \Gamma(\alpha,\beta)$$

be the conditional density derived from the SMC' transition probabilities, given that $X_i|Y_1,\dots,Y_i \sim \Gamma(\alpha,\beta)$. Note that with a small recombination rate, $p_{\alpha,\beta}$ is very close to $f_{\alpha,\beta}$. Then we can get:

$$\begin{aligned}
&\big(p_{\alpha,\beta}(t) - f_{\alpha,\beta}(t)\big)/\rho \\
=& e^{-t}\cdot\frac{(t\beta)^\alpha}{\Gamma(\alpha+1)}\left(M(\alpha,\alpha+1,-(\beta-1)t) - M(\alpha,\alpha+1,-(\beta+1)t)\right) \\
+& \frac{2t+e^{-2t}-1}{2}\cdot f_{\alpha,\beta}(t) + (1-e^{-2t})\cdot\frac{\Gamma(\alpha,\beta t)}{\Gamma(\alpha)} - 2t\cdot f_{\alpha,\beta}(t)
\end{aligned}$$

up to first order of $\rho$, where $M$ is Kummer's confluent hypergeometric function, also denoted ${}_1F_1$. The derivation is given in Supplement A.5.

*PDE approach to gamma approximation.* After obtaining a closed form, we would like to approximate it with a gamma distribution. Taking partial derivatives,

$$\frac{\partial f_{\alpha,\beta}(x)}{\partial \alpha} = f_{\alpha,\beta}(x) \cdot (-\psi^{(0)}(\alpha) + \log(\beta) + \log(x))$$
$$\frac{\partial f_{\alpha,\beta}(x)}{\partial \beta} = f_{\alpha,\beta}(x) \cdot (\alpha/\beta - x)$$

where $\psi^{(0)}$ is the digamma function. We can then find the coefficients $u, v$ such that

$$u \cdot \frac{\partial f_{\alpha,\beta}(x)}{\partial \alpha} + v \cdot \frac{\partial f_{\alpha,\beta}(x)}{\partial \beta} \approx (p - f)/\rho$$

Note that $u, v$ vary with $\alpha, \beta$. Then, we approximate

$$p_{\alpha,\beta}(x) \sim \mathit{\Gamma}(\alpha + \rho u, \beta + \rho v).$$

To find $u, v$, we evaluate the partial derivatives, as well at $(p - f)$, over a grid of numbers, and solve the least squares problem:

$$\arg\min_{u,v} ||u \cdot \frac{\partial f_{\alpha,\beta}(x)}{\partial \alpha} + v \cdot \frac{\partial f_{\alpha,\beta}(x)}{\partial \beta} + (p - f)/\rho||^2.$$

*Log-coordinates.* To better operate across scale, it is better to use coordinates specified in log-scale. If we use $\log_{10}(\alpha), \log_{10}(\beta)$ as our coordinates, by the chain rule,

$$\frac{\partial f_{\alpha,\beta}(x)}{\partial \log_{10}(\alpha)} = \frac{\partial f_{\alpha,\beta}(x)}{\partial \alpha} \cdot \frac{\partial \alpha}{\partial \log_{10}(\alpha)} = \frac{\partial f_{\alpha,\beta}(x)}{\partial \alpha} \cdot \frac{\alpha}{\log_{10}(e)}$$
$$\frac{\partial f_{\alpha,\beta}(x)}{\partial \log_{10}(\alpha)} = \frac{\partial f_{\alpha,\beta}(x)}{\partial \beta} \cdot \frac{\beta}{\log_{10}(e)}$$

Further, for improved interpretability and choice of grid boundaries, we wish to actually use the coordinates $\log_{10}(\mu), \log_{10}(C_v)$, where $\mu = \alpha/\beta$, $C_v = 1/\sqrt{\alpha}$. It follows that $\log_{10}(\mu) = \log_{10}(\alpha) - \log_{10}(\beta)$ and $\log_{10}(C_v) = -0.5 \cdot \log_{10}(\alpha)$. However, it turns out that those coordinates are not good for Taylor expansion, so that it is not true that perturbation of gamma in these coordinates is approximately equal to a linear combination of the partial derivatives. Instead, to get the respective change in $\log_{10}(\mu), \log_{10}(C_v)$, we use $\log_{10}(\alpha), \log_{10}(\beta)$ and, using linearity, simply write that

$$\mathit{\Delta} \log_{10}(\mu) = \mathit{\Delta} \log_{10}(\alpha) - \mathit{\Delta} \log_{10}(\beta)$$
$$\mathit{\Delta} \log_{10}(C_v) = -0.5\mathit{\Delta} \log_{10}(\alpha)\,.$$

*Emission step.* For this derivation, denote $T := X_{i+1}|\mathbf{Y}_{1:i}$. Suppose we know (or approximate) $p_{\alpha,\beta}$ as $X^{'}_{i+1} \sim \Gamma(\alpha', \beta')$. We use a Poisson emission model:

$$Y_{i+1}|(X_{i+1} = t) \sim Pois(2 \cdot \theta \cdot t)$$

Also, we assume an exponential prior for $X_{i+1}$. Then,

$$\begin{aligned}
\Pr(t|y_{i+1}) &= \Pr(y_{i+1}|t)\Pr(t)/\Pr(y_{i+1}) \\
&= (2\theta t)^{y_{i+1}} \cdot e^{-2\theta t} \cdot \frac{\beta^\alpha}{\Gamma(\alpha)} \cdot t^{\alpha-1} e^{-\beta t} / \Pr(y_{i+1}) \\
&\propto t^{\alpha+y_{i+1}-1} \cdot e^{-(2\theta+\beta)t}
\end{aligned}$$

and since this distribution must be normalized, we have that

$$\Pr(t|y_{i+1}) \sim \Gamma(\alpha + y_{i+1}, \beta + 2\theta).$$

That is:

$$X_{i+1}|\mathbf{Y}_{1:i+1} \sim \Gamma(\alpha, \beta + 2\theta)$$

if $Y_{i+1} = 0$ (is homozygous), and

$$X_{i+1}|\mathbf{Y}_{1:i+1} \sim \Gamma(\alpha + 1, \beta + 2\theta)$$

if it is heterozygous. If $Y_{i+1}$ is missing, we perform no updating.

**Combining forward and backward passes.** We have shown above that

$$\Pr(x_i|\mathbf{y}_{1:N}) \propto \hat{\alpha}(x_i) \cdot \int_0^\infty \Pr(x_i|x_{i+1}) \overleftarrow{\hat{\alpha}}(x_{i+1}) dx_{i+1} \cdot \frac{1}{\Pr(x_i)}.$$

We wish to combine gamma approximations from both the forward and backward passes to obtain a gamma approximation to the full posterior. First, we run the forward pass with the gamma approximation. This results in a gamma approximation to the forward density at each step, so that at position $i$ we have $\hat{\alpha}(x_i) \sim \Gamma(a, b)$.

Second, we run the forward pass with the gamma approximation on the reversed sequence, so that at position $i+1$ we have $\overleftarrow{\hat{\alpha}}(x_{i+1}) \sim \Gamma(a'', b'')$. Then, we apply the flow field once, to represent the change in uncertainty after taking a single transition step (from $i+1$ to $i$). This results in a new approximation, $\int_0^\infty \Pr(x_i|x_{i+1}) \overleftarrow{\hat{\alpha}}(x_{i+1}) dx_{i+1} \sim \Gamma(a', b')$. We then have, at each step $i$, a gamma approximation for both $X_i|\mathbf{Y}_{1:i}$ and $X_i|\mathbf{Y}_{i+1:N}$. Then we get

$$\begin{aligned}
\Pr(x_i|\mathbf{y}_{1:N}) &\propto \hat{\alpha}(x_i) \cdot \int_0^\infty \Pr(x_i|x_{i+1}) \overleftarrow{\hat{\alpha}}(x_{i+1}) dx_{i+1} \cdot \frac{1}{\Pr(x_i)} \\
&\propto x^{a-1} e^{-bx} \cdot x^{a'-1} e^{-b'x} / e^{-x} \\
&= x^{(a+a'-1)-1} e^{-(b+b'-1)}.
\end{aligned}$$

As this function must normalize to a proper density function, it follows that (based on the previous gamma approximations)

$$\Pr(x_i|\mathbf{y}_{1:N} \sim \Gamma(a + a' - 1, b + b' - 1)$$

So, to combine both steps, we simply sum the $\alpha$ and $\beta$ parameters, and subtract 1.

##### A.5 Gamma-SMC transition step derivation

The distribution of the next step is

$$\begin{aligned} p_{\alpha,\beta}(t) &= \int_{s=0}^{\infty} (\Pr(X_{i+1} = t|s, R_i)\Pr(R_i|s) + \Pr(X_{i+1} = t|s, \bar{R}_i)\Pr(\bar{R}_i|s))\Pr(X_i = s)\,ds \\ &\approx \int_{s=0}^{\infty} [q(t|s) \cdot 2\rho s + \delta(t-s) \cdot (1 - 2\rho s)]\, f_{\alpha,\beta}(s)\,ds \\ &= \int_{s=0}^{\infty} q(t|s) \cdot 2\rho s \cdot f_{\alpha,\beta}(s)\,ds + (1 - 2\rho t) f_{\alpha,\beta}(t) \end{aligned}$$

in first order of $\rho$. We get that the difference between consecutive positions (up to first order of $\rho$) is:

$$\begin{aligned} p_{\alpha,\beta}(t) - f_{\alpha,\beta}(t) &\approx \int_{s=0}^{\infty} q(t|s) \cdot 2\rho s \cdot f_{\alpha,\beta}(s)\,ds + (1 - 2\rho t) f_{\alpha,\beta}(t) - f_{\alpha,\beta}(t) \\ &= \rho \left[ \underbrace{\int_{s=0}^{\infty} q(t|s) \cdot 2s \cdot f_{\alpha,\beta}(s)\,ds}_{(*)} - 2t \cdot f_{\alpha,\beta}(t) \right] \end{aligned}$$

We proceed to develop $(*)$. We need to recall the following facts about the lower and upper incomplete gamma functions.

**Fact A1** *For all $t, \alpha, \beta > 0$,*

$$\begin{aligned} f_{\alpha,\beta}(x) &:= \frac{\beta^\alpha}{\Gamma(\alpha)} x^{\alpha-1} e^{-\beta x} \\ \gamma(s, x) &:= \int_0^x t^{s-1} e^{-t} dt \\ \Gamma(s, x) &:= \int_x^\infty t^{s-1} e^{-t} dt \\ \int_0^t f_{\alpha,\beta}(x) dx &= \frac{\gamma(\alpha, \beta t)}{\Gamma(\alpha)} \\ \int_t^\infty f_{\alpha,\beta}(x) dx &= \frac{\Gamma(\alpha, \beta t)}{\Gamma(\alpha)} \end{aligned}$$

**Fact A2** *For all $\beta$ (including non-positive) and for all $t, \alpha > 0$,*

$$\int_0^t x^{\alpha-1} e^{-\beta x} dx = \frac{t^\alpha}{\alpha} \cdot M(\alpha, \alpha+1, -\beta t),$$

*where $M$ is Kummer's confluent hypergeometric function, also denoted ${}_1F_1$. This generalizes the lower incomplete gamma function.*

We evaluate the integral separately at $s < t, s = t$ and $s > t$. For $s < t$,

$$\begin{aligned}\int_{s=0}^t q(t|s) \cdot 2s \cdot f_{\alpha,\beta}(s)\, ds &= \int_{s=0}^t \frac{e^{-(t-s)} - e^{-(t+s)}}{2s} \cdot 2s \cdot f_{\alpha,\beta}(s)\, ds \\ &= e^{-t} \int_{s=0}^t (e^s - e^{-s}) \cdot f_{\alpha,\beta}(s)\, ds \\ &= e^{-t} \cdot \frac{(t\beta)^\alpha}{\Gamma(\alpha+1)} \left(M(\alpha, \alpha+1, -(\beta-1)t) - M(\alpha, \alpha+1, -(\beta+1)t)\right)\end{aligned}$$

For $s = t$,

$$\begin{aligned}q(t|t) \cdot 2t \cdot f_{\alpha,\beta}(t)\, ds &= \frac{2t + e^{-2t} - 1}{4t} \cdot 2t \cdot f_{\alpha,\beta}(t)\, ds \\ &= \frac{2t + e^{-2t} - 1}{2} \cdot f_{\alpha,\beta}(t)\end{aligned}$$

For $s > t$,

$$\begin{aligned}\int_{s=t}^\infty q(t|s) \cdot 2s \cdot f_{\alpha,\beta}(s)\, ds &= \int_{s=t}^\infty \frac{1 - e^{-2t}}{2s} \cdot 2s \cdot f_{\alpha,\beta}(s)\, ds \\ &= \int_{s=t}^\infty (1 - e^{-2t}) \cdot f_{\alpha,\beta}(s)\, ds \\ &= (1 - e^{-2t}) \cdot \frac{\Gamma(\alpha, \beta t)}{\Gamma(\alpha)}\end{aligned}$$

To summarize,

$$\begin{aligned}&\left(p_{\alpha,\beta}(t) - f_{\alpha,\beta}(t)\right)/\rho \\ =& e^{-t} \cdot \frac{(t\beta)^\alpha}{\Gamma(\alpha+1)} \left(M(\alpha, \alpha+1, -(\beta-1)t) - M(\alpha, \alpha+1, -(\beta+1)t)\right) \\ +& \frac{2t + e^{-2t} - 1}{2} \cdot f_{\alpha,\beta}(t) + (1 - e^{-2t}) \cdot \frac{\Gamma(\alpha, \beta t)}{\Gamma(\alpha)} - 2t \cdot f_{\alpha,\beta}(t)\end{aligned}$$

up to first order of $\rho$.
